# PEC: a robust algorithm to reconcile pedigree and SNP-chip data on the basis of LD block, haplotype information, and Mendelian conflicts

**DOI:** 10.64898/2026.06.01.729286

**Authors:** Chuanke Fu, Quanshun Mei, Yuanxin Miao, Tao Xiang

## Abstract

**Motivation:** Pedigree errors frequently occur in livestock populations due to long-term manual record-keeping, which reduces the efficiency of breeding programs. Although several pedigree correction methods exist, their practical application is often limited by complicated procedures, high computational cost, and insufficient accuracy. Therefore, an effective and efficient solution for pedigree error correction is needed.

**Results:** We developed a new algorithm and software, PEC, to accurately and efficiently correct pedigree errors. The method matches haplotype fragments between candidate parents and offspring using estimated linkage disequilibrium patterns and subsequently checks for Mendelian conflicts to adjust the pedigree. Using simulated pig datasets, we compared PEC against SeekParentF90 and AlphaAssign in terms of accuracy, memory usage, and computation time. PEC demonstrated superior performance across all metrics. Furthermore, application of single-step genomic best linear unbiased prediction (ssGBLUP) in a real pig population showed that PEC corrected pedigrees significantly improved the accuracy and unbiasedness of genomic evaluations, highlighting the importance of pedigree error correction.

**Availability:** The PEC software is freely available at https://github.com/TXiang-lab/JPEC.

## Introduction

Pedigree is a key component throughout the development of modern livestock breeding. Since 1970s, when the best linear unbiased prediction (BLUP) method was established, pedigree information kept playing a crucial role in obtaining estimated breeding values (EBVs). In the era of genomic selection, the single-step genomic BLUP (ssGBLUP) method is used as a standard genomic evaluation method in routine livestock genomic evaluation. The ssGBLUP method relies on both pedigree and genomic information (Legarra et al. 2009, Christensen and Lund 2010, Christensen et al. 2012, Campos et al. 2018, Macedo et al. 2020, Abdollahi-Arpanahi et al. 2021), showing the importance of pedigree on genomic evaluation. Nevertheless, although pedigree plays an important role in animal breeding system, pedigree errors commonly exist in livestock populations. For instance, Visscher et al. (2002) reported that the pedigree error rate in UK dairy population ranged from 8.8% to 13.1%, whereas García-Ruiz et al. (2019) found that the pedigree error rate was 9.97% in the Mexican Holstein population. Such pedigree errors can be attributed to poor historical documentation or misrecordings, leading to a decrease in long-term genetic gain (Oliehoek and Bijma 2009). In human populations, kinship coefficients between individuals and the variance of kinship estimates across chromosomes may be informative to identify parent-offspring pairs, because human populations have limited background relatedness (Manichaikul et al. 2010, Hill and Weir 2011). However, this approach may be less effective in populations with the high background relatedness, small effective population size, and complex pedigree structure, such as livestock populations (Runge et al. 2022). In a livestock population, parent-offspring pairs and other close relatives may show similar kinship patterns, making them difficult to distinguish reliably using kinship coefficients. To identify and correct pedigree errors in populations with high background relatedness, several methods have been developed. At early stage, studies focused on using microsatellite markers or low-density single nucleotide polymorphism (SNP) array to verify the parentage (Riester et al. 2009). Later, with the rapid development of high-throughput sequencing technology, subsequent studies increasingly applied large SNP dataset to verify the parentage. Sargolzaei et al. (2014) identified parent-offspring pairs using an overlapping sliding windows approach across the whole genome. An alternative method was developed by Aguilar et al. (2014), who developed a software SeekParentF90 for the parentage identification by using Mendelian conflicts analysis, following the idea in Wiggans et al. (2009). In addition, Whalen et al. (2019) proposed a probabilistic method to infer the parents’ genotypes through assessing the likelihood of offspring’s genotypes with genotyping-by-sequencing data. These tools have proven useful, but they have some limitations. SeekParentF90 is sensitive to the chosen thresholds. Under defaulted threshold, it cannot detect misrecordings between close relatives (e.g., father and grandfather). AlphaAssign is computationally slow when applied to a large dataset. These limitations of existing software highlight the need for a robust, efficient algorithm and software for universal pedigree error correction in livestock breeding. Thus, in this study, we developed a new algorithm and software for pedigree error correction (PEC), which applies a novel approach integrating information of linkage disequilibrium (LD) patterns, haplotypes, and Mendelian conflicts to correct errors in the pedigree. We first tested PEC on a simulated pig dataset with five pedigree error rates and three error types, comparing its performance with SeekParentF90 (Aguilar et al. 2014) and AlphaAssign (Whalen et al. 2019). Then, we assessed the impact of pedigree correction on genomic evaluations using a real Chinese Yorkshire pig population. Our results show that PEC is both robust and efficient in identifying and correcting pedigree errors, thereby enhancing the accuracy of livestock breeding programs.

## Methods

### Algorithm of PEC

The PEC algorithm operates in two sequential steps: first, matching haplotype fragments (LD blocks) between candidate parents and offspring based on estimated linkage disequilibrium (LD) patterns, and second, detecting Mendelian conflicts between these pairs to perform pedigree error correction.

Genetic information is transmitted from parent to offspring via haplotype inheritance. However, chromosomal crossover during prophase I of meiosis results in physical exchange of haplotype segments between homologous chromosomes. To account for this, we propose dividing haplotypes into smaller fragments, where each fragment is defined as a fundamental unit of genetic transmission. These fragments allow precise tracking of inherited genetic information, even when recombination occurs. Larger genomic segments can then be reconstructed by aggregating contiguous fragments. A fragment is equivalent to a linkage disequilibrium (LD) block, based on the principle that adjacent SNPs within an LD block exhibit stronger allelic association than SNPs outside the block (Cardon and Abecasis 2003). The size and distribution of LD blocks are determined using the algorithm described by Gabriel et al. (2002). A LD block was defined as a region in which fewer than 5% of informative SNP pairs show strong evidence of historical recombination. Since both offspring and candidate parents are diploid, one of the offspring’s two haplotypes must originate from the parent. To determine the transmitted haplotype, the following steps are performed:

1. Label LD blocks sequentially (e.g., LD block 1, LD block 2) across the entire haplotype
2. For each LD block (e.g., LD block 1), compare the offspring’s haplotypes (haplotypes 1 and 2) to the candidate parent’s haplotypes (haplotypes 3 and 4). If all SNPs in the offspring’s haplotype 1 match either haplotype 3 or 4 of the parent within the LD block, assign a score of 1; otherwise, assign 0.
3. Repeat this comparison for all LD blocks on haplotype 1 and calculate the total matching score. Perform the same analysis for haplotype 2.
4. The haplotype with the higher matching score is identified as the transmitted haplotype, with the higher score designated as the final count.

The percentage of matching LD blocks between the offspring and candidate parent was calculated as an indicator to evaluate the parentage relationships. In addition, Mendelian conflicts—defined as opposing homozygous genotypes at a SNP between parent and offspring (Hayes 2011) — are incorporated in our algorithm. Even true parent-offspring pairs may exhibit a small number of Mendelian conflicts due to the genotyping errors. However, conflicts between true pairs are an order of magnitude lower than those involving non-parents (Hayes 2011). The number of Mendelian conflict SNPs is counted between an offspring and its candidate parent. To prioritize these conflicts, the percentage of unmatched LD blocks containing Mendelian conflict SNPs is subtracted from the parentage score, thereby weighting Mendelian conflicts more heavily than other mismatches.

The overall model of PEC can be described as:

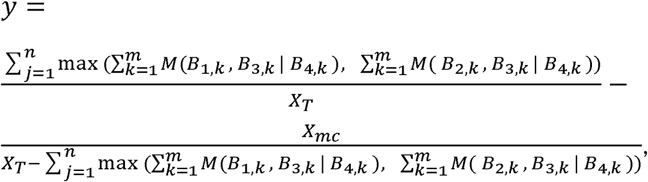

where y is the matching score between an offspring and its candidate parent; *B*_1,*k*_ and *B*_2,*k*_ are the k-th LD block on haplotype 1 and 2 for the offspring, while *B*_3,*k*_ and *B*_4,*k*_ are the k-th LD block on haplotype 3 and 4 for the candidate parent; the function *M*(*B*_1,*k*_ *, B*_3,*k*_ *| B*_4,*k*_) returns 1 if all the SNPs on *B*_1,*k*_ completely match the SNPs on either *B*_3,*k*_ or *B*_4,*k*_, otherwise, it returns 0. Similarly, *M*(*B*_2,*k*_ *, B*_3,*k*_ | *B*_4,*k*_) evaluates haplotype 2 of the offspring against the candidate parent’s haplotypes; k ranges from 1 to m, and m is the total number of LD blocks on chromosome j; the algorithm calculates two sums: 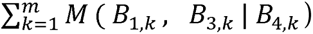 and 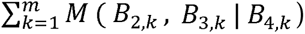, and the higher of these two sums is selected as the final count for chromosome j. This process is repeated across all chromosomes (j ranges from1 to n, where n is the total number of chromosomes). *X_T_* is the total number of LD blocks identified by a phasing software; *X_mc_* is the number of LD blocks containing Mendelian conflict SNPs. The percentage of Mendelian conflict-containing LD blocks among non-matching LD blocks is subtracted from the parentage relationship score to prioritize Mendelian conflicts over other mismatches.

We use father identification as an example to illustrate the method of identifying a parent of an individual in pedigree. Both the father and the individual must be genotyped. The pedigree file needs to include five columns: individual ID, father ID (could be wrong), mother ID (could be wrong), birth date, and gender. Identification is performed among all the genotyped males whose age exceeds the individual’s birth date by at least the minimum recorded father-offspring age gap in the pedigree.

PEC is implemented in Julia, a high-performance language for scientific computing (Bezanson et al. 2017). User’s Guide is demonstrated in the github site https://github.com/TXiang-lab/JPEC.

### Simulated data

Phenotypic and genotypic data sets were simulated by the software QMSim version 1.10 (Sargolzaei and Schenkel 2009). The population mimicked a pig selection scheme, with a trait was only motivated by additive genetic effects. The process of simulation followed by Xiang et al. (2018). The simulation included a historical population undergoing mutation and drift, followed by two recent populations under selection. In the historical population, the effective population size decreased from 500 to 65 and then increased to 220 in the final generation, from which 20 males and 200 females were sampled as founders of the first recent population. Individuals in the recent populations were selected to breed the next generation based on high estimated breeding values (EBVs) predicted by BLUP. The second recent population was founded by 100 males and 500 females selected from the first recent population and then underwent 13 generations of selection.

The simulation included 18 autosomes, each 120 cM in length. Each autosome contained 200 randomly distributed biallelic quantitative trait loci (QTLs) and 13,000 biallelic markers. The simulated population generated pedigree records, phenotypic records, and genotypic records for all individuals. A real pig population usually has the limitation that the pedigree records are typically the most complete, followed by phenotypic records, and genotypic records the least. Therefore, we retained pedigree records for all 74,134 individuals across generations 1 to 10, phenotypic records for all 36,634 individuals in generations 6 to 10, genotypic records for 50% of individuals in each generation from 6 to 10 (18,315 individuals), randomly selected under the condition that their sires were genotyped. To simulate the usage of genotypes in a real pig population, genotypes were used after quality control. SNPs were removed with minor allele frequency (MAF) lower than 0.05, or a significant deviation from Hardy Weinberg equilibrium (p < 10e-7), which is most likely due to technical artifacts. After quality control, 33,547 SNPs of 18,315 pigs were retained.

To simulate varying errors in pedigree, we created nine scenarios, combining three types (random, full-sib, and sire errors) across three rates (1%, 5%, and 10%) of pedigree errors (Table 1). Three types of error records were focused on the male pigs: Random error, where pigs’ fathers were replaced by another random male pig; Full-sib error, where pigs’ fathers were replaced by their fathers’ full-siblings (uncles); and Sire error, where pigs’ fathers were replaced by their grandfathers in the pedigree. The pedigree error rates were calculated as the proportion of pigs with incorrect fathers to all genotyped pigs, with rates of 1%, 5%, and 10% in the simulations.

**Table 1.** Scenarios across three types and three rates of pedigree errors in the simulation.

| Scenario | Type of Error | Rate of Errors |
| --- | --- | --- |
| 1 | Random | 1% |
| 2 | Random | 5% |
| 3 | Random | 10% |
| 4 | Full-sib | 1% |
| 5 | Full-sib | 5% |
| 6 | Full-sib | 10% |
| 7 | Sire | 1% |
| 8 | Sire | 5% |
| 9 | Sire | 10% |

### Real data

#### Phenotypic records

Phenotypic records were collected from a Chinese Yorkshire (CY) pig population, including measurements of average daily gain (ADG), backfat thickness (BF), loin muscle depth (LMD), and age to 100 kg (AGE). ADG was calculated from birth to the off-test date. Records for ADG, BF, LMD, and AGE were available for 33,579, 33,601, 33,587, and 33,681 pigs, respectively, between 2013 and 2021. The pedigree spanned January 1, 2013, to August 29, 2021, and included 375,636 individuals.

#### Genotypes

In the population, 17,488 animals were genotyped, where 10,281 were in the Illumina PorcineSNP60 Genotyping BeadChip and 7,207 were in Zhongxin NO.1 chip. SNPs were mapped to chromosomes based on the pig genome build 10.2 (Groenen et al. 2012). First, the two kinds of chips were imputed to the same size as 40,348 SNPs and combined following our previous methods (Mei et al. 2022). Specifically, SNPs shared between the two kinds of chips were used as a backbone, and the remaining SNPs were then imputed to bring genotypes from one chip up to the marker density of the other. The same principle of merging chip datasets and imputing missing markers was adopted to generate a consistent genotype panel for downstream analyses. After quality control using the same criteria as in the simulation above, 32,234 SNPs of 17,488 pigs remained.

#### Evaluation of pedigree correction accuracy

Pedigree correction accuracy was calculated as the proportion of incorrectly assigned fathers successfully corrected in simulations. LD blocks utilized in PEC were derived from LD block set data and haplotype information. Plink v1.9 was recommended for generating LD block sets (Purcell et al. 2007). Variant IDs and position information are recorded for each block. The default maximum length of LD blocks is 200 KB. However, a study by Amaral et al. (2008) shows that LD block can extent up to 400 KB in some European breeds. To explore the effect of maximum lengths of LD block on PEC, three maximum lengths were set as 200 KB, 500 KB, and 1000 KB. Beagle 5.2 was recommended for haplotype phasing by Miar et al. (Miar et al. 2017). Beagle 5.2 uses a hidden Markov model along the chromosome to infer each individual’s haplotypes (Browning et al. 2021). Alternative software may also be used for generating LD block set data and haplotype inference. Therefore, the running time and RAM usage of these steps may vary depending on the software, parameter settings, and number of threads used.

We compared accuracies, memory usage, and computation time between PEC with two other existing methods of pedigree correction: SeekParentF90 v1.56 (Aguilar et al. 2014;), which seeks parents based on Mendelian conflicts, and AlphaAssign (Whalen et al. 2019; executable SHA256 checksum: ab42f65ed295b49d08f5af81e7f6b1c404e0735b639bd1be9a35acd4f96ee7f6), which uses a probabilistic method for pedigree correction. Both PEC and SeekParentF90 support genome-wide parent identification either with or without pre-specifying the candidate individuals. On the contrary, AlphaAssign must pre-specify the candidate individuals before operation. All nine scenarios (3 error types × 3 error rates) of pedigree errors were evaluated. To ensure comparable accuracies, memory usage, and computation time across the three software, these individuals having incorrect parent recordings were provided to AlphaAssign, PEC, and SeekParentF90.

In addition, AlphaAssign and SeekParentF90 do not support multithread, but PEC can work in the multithread environment. Thus, the memory usage and computation time of running PEC with 1, 5, 10, and 20 threads were compared to the other two software. The memory usage was measured as the maximum resident memory in the process of pedigree correction. Accuracies of pedigree correction were evaluated in all the nine scenarios. Memory usage and computation time were only evaluated in the scenarios of the Random type because comparisons on memory usage and computation time are not influenced by the types of pedigree errors.

#### Statistical models

Univariate animal models were applied for genomic evaluations in both simulated data and real data. The statistical model for the trait in the simulated data was as follows:

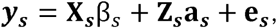

where ***y_s_*** is the vector of phenotypes; β*_s_* is the vector of fixed effects, which include the mean of the trait and sex effect; ***a_s_*** is a vector of additive genetic effect; ***e_s_*** is a vector of random residual effect. ***X_s_*** and ***Z_s_*** are the respective incidence matrices. In real data, ADG, BF, and LMD were analysed in the same statistical models, as follows:

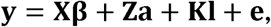

where *y* represents vectors of phenotypic records for ADG, BF, and LMD in the statistical models, respectively; **β** is a vector of fixed effects, which include sex effect, test section effect, herd-year effect, and covariate on off-test age, defined as the number of days from birth to the off-test date of an individual; **a** is a vector of the additive genetic effects for ADG, BF, and LMD, respectively; **l** is a vector of random litter effects; **X, Z,** and **K** are the corresponding incidence matrices; **e** is the vector of random residual effects. Random litter effects were assumed to follow a normal distribution as 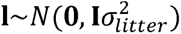, where **I** is an identity matrix, 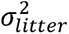 is the variance of litter effect. Random additive genetic effects **a** were assumed to follow normal distribution as 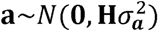, where **H** is the combined genomic and pedigree-based relationship matrix relationship matrix as defined in Christensen et al. (2012), computed as

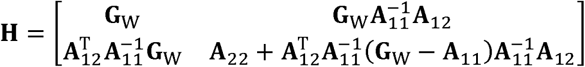

where **G**_W_ is the adjusted genomic relationship matrix for genotyped animals. Matrices **A**_11_, **A**_22_, and **A**_12_ are pedigree relationship matrices among genotyped, among non-genotyped and between genotyped and non-genotyped animals, respectively. 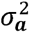 is the additive genetic variance.

The statistical model of AGE was similar to the models of the other three traits, except that the covariate was on off-test weight instead of off-test age.

#### Genomic evaluations

To investigate the effectiveness of the methods of pedigree correction on genomic evaluations, single-step genomic evaluations were implemented with either an original incorrect pedigree or a pedigree corrected by PEC, Seekparentf90, and AlphaAssign in both simulated data and real data. The inverse of the combined genomic and pedigree-based relationship matrix (**H**^−1^) was constructed as defined in Christensen and Lund (2010) and Christensen et al. (2012). Note that weighting factor ω was fixed at 0.05 for all the analyses in this study. The performances of genomic evaluations were then evaluated using software DMU v5.3 (Madsen and Jensen 2013).

#### Genomic evaluations of simulated data

In simulated data, evaluation accuracy was measured as correlation coefficients between true breeding values (TBV) and EBVs in validation set. Meanwhile, regression coefficient of TBV on EBV was calculated as an indicator of dispersion bias of the genomic evaluation. A greater absolute deviation of the regression coefficient from 1 indicates a stronger bias. EBVs were obtained using the preconditioned conjugate gradient method in DMU software (Madsen and Jensen 2013).

Simulated dataset was split into training and validation sets based on the generations: individuals in generation 6-9 were set as training set and individuals in generation 10 were set as validation set. Thus, training set had 29401 individuals, and among them 14699 were genotyped; validation set had 7233 individuals, and among them 3616 were genotyped. A Hotelling-Williams t-test at a 5% confidence level was performed to evaluate the statistically significant differences in evaluation accuracy between scenarios of different rates and types of pedigree error. The results were compared between different methods.

#### Genomic evaluations of real data

Similar to simulated dataset, based on the cutting-off date, phenotypic records were divided into training set and validation set. Individuals were born before August 1st, 2020 were put into training set and the rest individuals were put into validation set. Evaluation accuracy and regression coefficient were analysed. The number of pigs with records in reference and validation sets was shown in Table 2.

**Table 2.** The number of pigs with records for average daily gain (ADG), backfat (BF), lion depth (LD), and age to 100 kg (AGE) in the training and validation groups.

| Trait | Training | Validation |
| --- | --- | --- |
| ADG | 26442 | 7137 |
| BF | 26467 | 7134 |
| LD | 26452 | 7135 |
| AGE | 26544 | 7137 |

ADG average daily gain, BF backfat, LD lion depth, AGE age to 100 kg

In the real Chinese Yorkshire pig population, evaluation accuracy were determined as correlation between corrected phenotypes (**Y***_c_*) and EBV (**â**) in validation populations (*cor* (**Y**_c_, **â**)), following Christensen et al. (2012). Corrected phenotypes were equal to the sum of estimated additive genetic effects and residual effects for each individual (**Y**_c_ = **â** + **ê**). In addition, regression coefficients of **Y***_c_* on EBV were calculated as an indicator of dispersion bias of the genomic evaluation for each trait. We conducted a Hotelling-Williams t-test at a 5% confidence level to assess the statistically significant differences in evaluation accuracy between scenarios involving the original pedigree and pedigree corrected by the different algorithms (PEC, SeekParentF90, AlphaAssign).

## Results

### Evaluation of pedigree correction accuracy

LD block maximum lengths (200KB, 500KB, and 1000KB) had little impact on the performances of pedigree correction in PEC, thus we showed results of pedigree correction based on 200KB-LD block. Although all the three methods showed high accuracies (>97%) in pedigree error correction, PEC outperformed the other two methods in terms of the accuracies of pedigree correction, regardless of types of pedigree errors (Fig. 2a, 2b, and 2c). For all the different types and rates of errors, PEC always achieved 100% accuracy, while SeekParentF90 and AlphaAssign showed slightly lower accuracies: in the type of Random error, both PEC and SeekParentF90 showed 100% accuracies, while AlphaAssign showed marginally lower accuracy across all rates of pedigree errors; in the type of Full-sib error, AlphaAssign outperformed or equalled the accuracy of SeekParentF90 (99.5%) across all rates of pedigree errors; in the type of Sire error, AlphaAssign (≥ 99.3%) outperformed SeekParentF90 (≥ 97.8%) across all rates of pedigree errors.

**Fig. 1.**
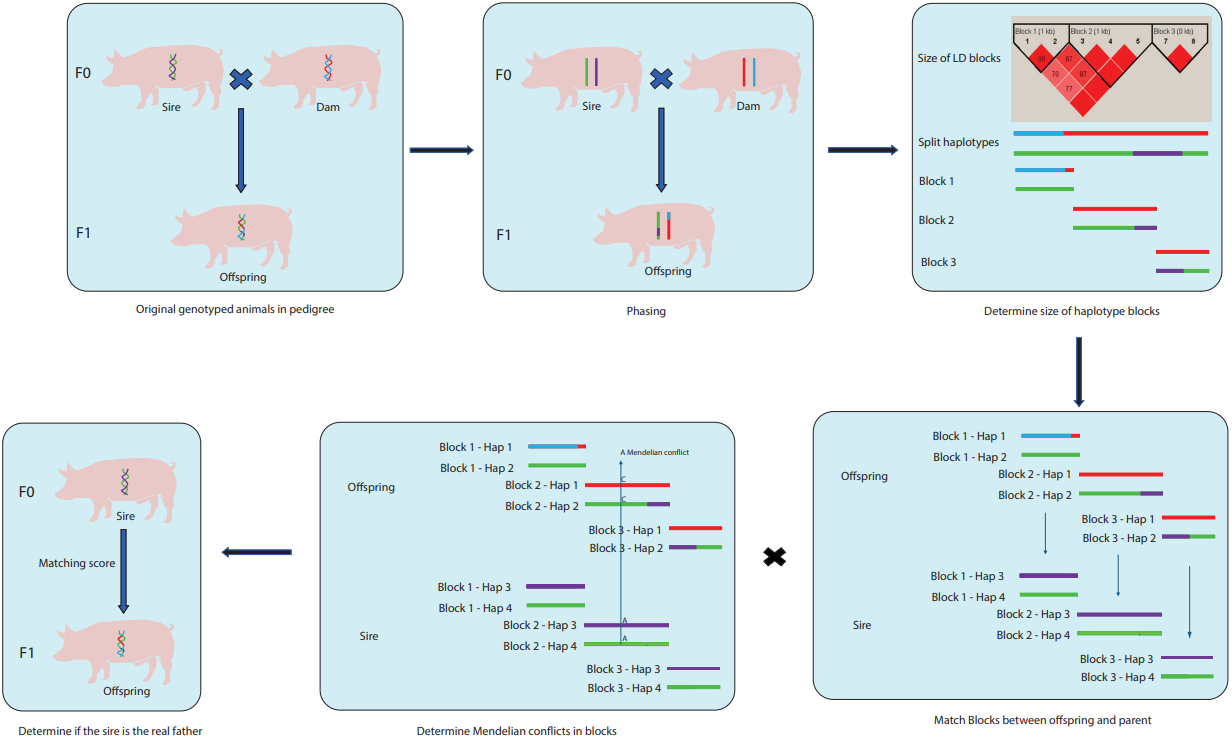
Overview of PEC workflow

**Fig. 2.**
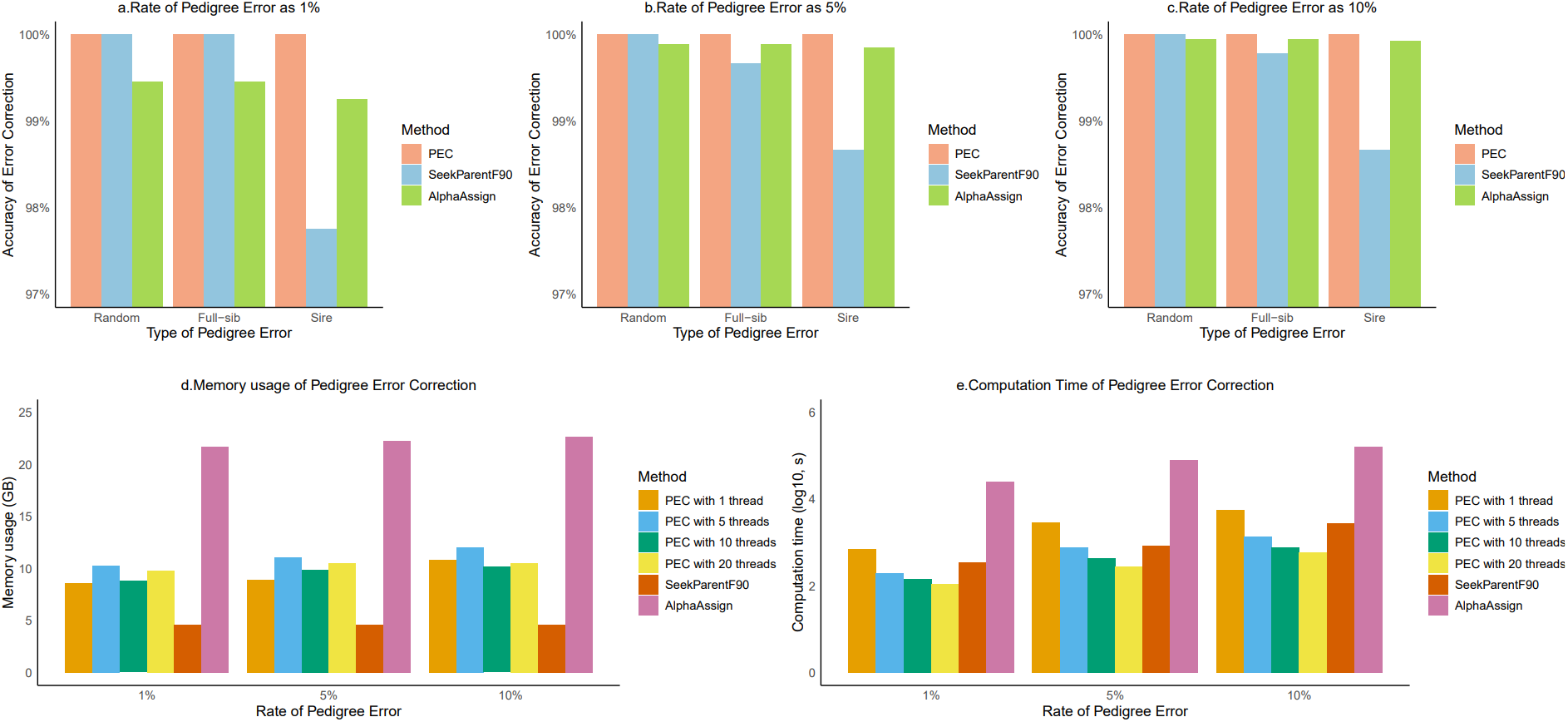
Comparison of accuracy, memory usage, and computation time across different types and rates of pedigree errors

For all the three software (PEC, AlphaAssign and SeekParentF90), memory consumption remained stable across different scenarios (Fig. 2d). Among them, SeekParentF90 required the least memory, staying constant as 4.6 GB. PEC required about 10 GB regardless of the number of threads. AlphaAssign required the most memory, reaching to about 21 GB.

The computation time increased as the rate of pedigree error increased from 1% (183 individuals) to 10% (183 individuals) (Fig. 2e). For PEC, the computation time decreased considerably as the number of used threads increased. When 20 threads were used in PEC, the computation time shrunk to the minimum, approaching to 110 seconds for an error rate of 1%, 284 seconds for a rate of 5%, and 584 seconds for a rate of 10%. The computation time of SeekParentF90 similar to PEC with 1 to 5 threads, reaching to 343 seconds for a rate of 1%, 854 seconds for a rate of 5%, 854, and 2735 seconds for a rate of 10%. For AlphaAssign, the computation time was the longest among all the three software, up to 25916 seconds for a rate of 1%, 81489 seconds for a rate of 5%, 160729 seconds for a rate of 10%.

### Genomic evaluations

#### □D. Genomic evaluations of simulated data

Accuracies and biasedness of genomic evaluations declined gradually as pedigree error rates increased (Table 3). For Random errors, evaluation accuracy decreased from 0.608 to 0.597, and biasedness decreased from 0.880 to 0.831 as error rates rose from 1% to 10%. Full-sib and Sire errors showed similar but less pronounced trends compared to Random errors.

**Table 3.** Accuracies and regression coefficients of genomic evaluation for the simulated trait across three rates and three types of pedigree error.

| Rate | Type | Original |  | PEC |  | SeekParentF90 |  | AlphaAssign |  |
| --- | --- | --- | --- | --- | --- | --- | --- | --- | --- |
|  |  | Accuracy | Reg | Accuracy | Reg | Accuracy | Reg | Accuracy | Reg |
| 1% | Random | 0.608<sup>a</sup> | 0.880 | 0.610<sup>c</sup> | 0.888 | 0.610<sup>c</sup> | 0.888 | 0.610<sup>b</sup> | 0.888 |
|  | Fullsib | 0.609<sup>a</sup> | 0.885 | 0.610<sup>c</sup> | 0.888 | 0.610<sup>c</sup> | 0.888 | 0.610<sup>b</sup> | 0.888 |
|  | Sire | 0.609<sup>a</sup> | 0.886 | 0.610<sup>c</sup> | 0.887 | 0.610<sup>b</sup> | 0.887 | 0.610<sup>b</sup> | 0.887 |
| 5% | Random | 0.604<sup>a</sup> | 0.860 | 0.610<sup>c</sup> | 0.888 | 0.610<sup>c</sup> | 0.888 | 0.610<sup>b</sup> | 0.888 |
|  | Fullsib | 0.608<sup>a</sup> | 0.879 | 0.610<sup>c</sup> | 0.888 | 0.610<sup>b</sup> | 0.888 | 0.610<sup>b</sup> | 0.888 |
|  | Sire | 0.609<sup>a</sup> | 0.883 | 0.610<sup>c</sup> | 0.886 | 0.610<sup>bc</sup> | 0.886 | 0.610<sup>b</sup> | 0.886 |
| 10% | Random | 0.597<sup>a</sup> | 0.831 | 0.610<sup>c</sup> | 0.888 | 0.610<sup>c</sup> | 0.888 | 0.610<sup>b</sup> | 0.888 |
|  | Fullsib | 0.606<sup>a</sup> | 0.872 | 0.610<sup>c</sup> | 0.888 | 0.610<sup>c</sup> | 0.888 | 0.610<sup>b</sup> | 0.888 |
|  | Sire | 0.608<sup>a</sup> | 0.876 | 0.609<sup>d</sup> | 0.884 | 0.609<sup>b</sup> | 0.883 | 0.609<sup>c</sup> | 0.884 |

Here accuracies and regression coefficients are calculated with either an original incorrect pedigree in whole validation population as scenario Original or a pedigree corrected of PEC, SeekParentF90, and AlphaAssign, respectively
Reg: regression coefficient, which of TBV on EBV were calculated as an indicator of dispersion bias of the genomic evaluation
For accuracy, different superscripts of small letters among scenarios indicate significant differences ( $p < 0.05$ ) by Hotelling-Williams t-test

All the three pedigree-correction software improved evaluation accuracy and reduced biasedness across different error types (Table 3). For the type of Random and Full-sib errors, evaluation accuracy reached a maximum of 0.610, and regression coefficient reached a maximum of 0.888. In the type of Sire error, corrected pedigree showed a smaller improvement of accuracy when compared with the other two types of error, with evaluation accuracy ranging from 0.609 to 0.610 and biasedness ranging from 0.883 to 0.887. Moreover, Hotelling-Williams t-test indicated that PEC significantly outperformed (p-value<0.05) other two software, followed by SeekParentF90, and AlphaAssign was ranked third. PEC outperformed AlphaAssign in all the different scenarios while PEC surpassed SeekParentF90 in Sire errors (1% and 10% rates) and Full-sib errors (5% rate). In other scenarios, PEC performed similarly to SeekParentF90.

#### □D. Genomic evaluations of real data

Based on findings from simulated data, LD blocks were constructed using the default maximum length of 200 kb for pedigree correction with PEC in the Chinese Yorkshire population. Evaluation accuracies and biasedness were compared between uncorrected pedigrees and pedigrees corrected by PEC, SeekParentF90, and AlphaAssign for four traits: ADG, BF, LMD, and AGE (Table 4). Overall, in terms of evaluation accuracies, using PEC to correct pedigree did not significantly improve evaluation accuracy for ADG, BF, and LMD, but significantly improved evaluation accuracy for AGE (evaluation accuracy increased from 0.14 before pedigree correction to 0.142). However, evaluation accuracies of any traits did not increase when using SeekParentF90 or AlphaAssign to correct pedigree.

**Table 4.** Accuracies and regression coefficients of genomic evaluation for average daily gain (ADG), backfat (BF), lion depth (LD), and age to 100 kg (AGE) in CY population.

| Trait | Original |  | PEC |  | SeekParentF90 |  | AlphaAssign |  |
| --- | --- | --- | --- | --- | --- | --- | --- | --- |
|  | Accuracy | Reg | Accuracy | Reg | Accuracy | Reg | Accuracy | Reg |
| ADG | 0.276<sup>a</sup> | 0.872 | 0.275<sup>a</sup> | 0.880 | 0.276<sup>a</sup> | 0.873 | 0.275<sup>a</sup> | 0.874 |
| BF | 0.348<sup>b</sup> | 0.891 | 0.348<sup>ab</sup> | 0.894 | 0.348<sup>a</sup> | 0.889 | 0.348<sup>ab</sup> | 0.890 |
| LMD | 0.235<sup>ab</sup> | 0.960 | 0.236<sup>b</sup> | 0.966 | 0.236<sup>ab</sup> | 0.961 | 0.235<sup>a</sup> | 0.960 |
| AGE | 0.140<sup>a</sup> | 0.609 | 0.142<sup>b</sup> | 0.634 | 0.140<sup>a</sup> | 0.614 | 0.140<sup>a</sup> | 0.614 |

Here accuracy and regression coefficients are calculated with either an original pedigree as scenario Original or a pedigree corrected of PEC, SeekParentF90, and AlphaAssign, respectively.
Reg: regression coefficient, which of corrected phenotypes on EBV were calculated as an indicator of dispersion bias of the genomic evaluation
For accuracy, different superscripts of small letters among scenarios indicate significant differences ( $p < 0.05$ ) by Hotelling-Williams t-test

In terms of biasedness, pedigree correction decreased biasedness for all the four traits, and PEC outperformed SeekParentF90 and AlphaAssign. More specifically, when using PEC to correct pedigree, it reduced biasedness for all the four traits. However, biasedness was only reduced for LMD and AGE when using SeekParentF90 to correct pedigree, while AlphaAssign only reduced biasedness for ADG and AGE. Furthermore, PEC, SeekParentF90, and AlphaAssign performed differently in improving biasedness. For instance, using PEC to correct pedigree for AGE showed the greatest improvement of biasedness, achieving a 0.025 decreased biasedness.

## Discussion

To determine and correct error recordings in pedigree, we developed a robust method, PEC, to reconcile pedigree and SNP-chip data based on LD block, haplotype information, and Mendelian conflicts. In simulation, PEC demonstrated high correction accuracy (>97%), stable memory usage (∼10 GB), and scalable computational efficiency (110–584 seconds with multithreading). In a real pig population, pedigree correction by PEC reduced bias in genomic evaluation. In general, PEC showed superior performance in both correction accuracy and bias reduction in genomic evaluation compared to existing two methods SeekparentF90 and AlphaAssign.

To determine the impact of maximum length to construct LD blocks, we examined three different maximum lengths including 200KB, 500KB, and 1000KB. The results showed that although the number of matched fragments and the matching scores varied with different maximum lengths used to construct LD blocks, the ranking remained unchanged. Therefore, the three different maximum lengths did not affect the performance of PEC. It can be attributed to two reasons. First, we used Beagle5.2 for haplotype phasing before constructing LD blocks. The high phasing accuracy (>90%) ensured reliable transmission tracking. Second, we mimicked a sophisticated pig selection scheme, existing the stable long haplotype blocks. When we set the maximum length of constructed LD block as 1000KB, the stable long haplotype block sizes were within a LD block, which can be regarded as the genetic units transmitted to the next generation. When we set the maximum length of constructed LD block as the default 200KB, the stable long haplotype block was fragmented into several short haplotype blocks, and these short haplotype blocks can still be accurately matched between parent and offspring.

Using SeekParentF90 to correct pedigree performed worse in the type of Sire error than in the types of Random error and Full-sib error (Fig. 2a, 2b, and 2c). It can be attributed to the high genotype similarity between true and erroneous sires.

SeekParentF90 corrects pedigree by identifying Mendelian conflicts between parents and offspring, using a default threshold of 1% of the total number of SNPs (Wiggans et al. 2009). For some parent-offspring pairs in the type of Sire error, the number of Mendelian conflicts between wrong sires and offspring was lower than the threshold, making identification of parent more challenging.

In the simulation, both evaluation accuracy and regression coefficient declined with increasing rate of pedigree error (Table 3). These results indicate that pedigree errors did negatively impact genomic evaluation, consistence with previous studies (Roughsedge et al. 2001, García-Ruiz et al. 2019). PEC mitigated these effects, outperforming AlphaAssign across all error types and rates. Statistical significance (Hotelling-Williams t-test, p < 0.05) confirmed subtle differences in error distributions between different pedigree correction methods.

In the real pig population, using PEC to correct pedigree siginificantly improved evaluation accuracy for AGE, rather than ADG, BF, and LMD (Table 4). The results suggest that a larger proportion of genetic variation for AGE may be captured by pedigree information rather than by SNPs in ssGBLUP, in contrast to ADG, BF, or LMD. Using SeekParentF90 and AlphaAssign to correct pedigree did not siginificantly improve evaluation accuracy for any of the four traits. It is likely attributed to the default thousholds to select parents in SeekParentF90 and AlphaAssign, which are relatively tolerant and may not effectively address potential issues, such as sire errors. However, using SeekParentF90 or AlphaAssign can still effectively correct other types of pedigree error, such as Random error, thereby reducing biasness observed in genomic evaluation after pedigree correction through PEC, SeekParentF90, and AlphaAssign. Furthermore, pedigree correction showed the greatest improvement for AGE compared to ADG, BF, or LMD. The results also suggest that a larger proportion of genetic variation for AGE may be captured by pedigree information rather than by SNPs, in contrast to ADG, BF, or LMD.

In a real population, PEC, SeekParentF90, and AlphaAssign were applied to all genotyped individuals to correct pedigree. However, these methods were applied to individuals having incorrect parent recordings in each scenario to evaluate pedigree correction accuracy. To further assess their performance, we applied PEC, SeekParentF90, and AlphaAssign to all genotyped individuals in simulation.

As a result, PEC identified and corrected all genotyped individuals. PEC required memory as about 17.7 GB, and computation time as about 20.6 hours with a thread, which was reduced to about 30 minutes with 64 threads. SeekParentF90 performed the same with all genotyped individuals compared to with individuals having incorrect parent recordings in each scenario. In contrast, AlphaAssign performed the worst: it failed to identify the true fathers of eight individuals, required the largest memory as about 27.7 GB, and took about 16 days.

From a practical perspective, computation time for PEC should be considered for both preprocessing and pedigree correction. Previous benchmarking study has shown that haplotype phasing in large datasets may require substantial computation time and memory(De Marino et al. 2022). De Marino et al. (2022) reported that phasing a whole-genome sequencing dataset with 2280 individuals and 1 million variants required about 11.2 h of CPU time and 19 GB of memory. The runtime of PEC itself is approximately proportional to the number of comparisons between offspring and candidate parents and to the amount of LD-block comparison, while decreasing with the number of threads used. Our results showed that PEC scales well with multithreading while maintaining stable memory usage. For example, in our simulation dataset containing 18,315 genotyped pigs and 33,547 SNPs, PEC required about 30 min and 17.7 GB of memory when multiple threads were used.

In the simulation, SNPs were complete and filtered with MAF>0.05. However, filtering SNPs with MAF>0.01 is also common in genomic evaluation. After long term selection, some SNPs linked to causal variants may become nearly fixed, resulting in low-MAF SNPs. Low-MAF SNPs make phasing harder because the algorithm of phasing for Beagle relies on identifying haplotypes shared across individuals (Miar et al. 2017). A SNP with low MAF indicates only limited individuals carry the SNP, which limits haplotype sharing. Furthermore, the LD between low-MAF SNPs and their neighbouring SNPs may be weak. In addition, SNPs genotyping can include sporadic missing calls and genotyping errors in practice. These issues can also reduce phasing accuracy, thus affect the performance of PEC. To assess the impact of low-MAF SNPs, or sporadic missing SNPs and genotyping errors on pedigree correction, we evaluated pedigree correction accuracy using two modified genotype datasets: (1) a MAF of 0.01, or (2) 1% of SNPs randomly sporadic missing and 0.5% of SNPs randomly containing error across all individuals. For both datasets, PEC outperformed SeekParentF90 and AlphaAssign, with pedigree correction accuracy reaching 100% in all scenarios (Table S1 and S2). The consistent performance indicated the robustness and effectiveness of PEC.

To maximize the detection of potential parents, we included gender information for genotyped individuals in pedigree when running PEC. However, a traditional pedigree format with four columns (ID, Sire, Dam, Birthday) can also be used for pedigree correction. Although the scale of potential parents is determined in sires and dams rather than all genotyped individuals in the case, the running speed is improved with possibly only a slight compromise in correction accuracy.

In general, PEC is a robust method to reconcile pedigree and SNP-chip data based on LD block, haplotype information, and Mendelian conflicts. PEC is highly recommended for pedigree correction due to its superior performance in both simulation and real population. Comprehensively considering the accuracy, memory usage, and computation time, PEC is highly recommended for routine pedigree correction in pig breeding, offering a practical solution to enhance the reliability of genomic evaluations.

## Data Availability Statement

The datasets in the current study are available from the corresponding author on reasonable request.

## Acknowledgements

Not applicable.

## Authors’ contributions

TX, CF, and QM conceived the study. QM and CF developed the PEC software. CF, QM, and YM carried out the data analysis. CF drafted the manuscript, TX and QM revised the manuscript. TX provided the data. All authors read and approved the final version. CF and QM contributed equally to this work.

