## Supplementary Material for "PEC: a robust algorithm to reconcile pedigree and SNP-chip data on the basis of LD block, haplotype information, and Mendelian conflicts"

**Supplementary Table S1 accuracy of pedigree correction of genotypes with a MAF as 0.01 (%)**

| Rate | Type | PEC | SeekParentF90 | | AlphaAssign | |
| --- | --- | --- | --- | --- | --- | --- |
| 1% | Random | 100 | | 100 | | 99.5 |
|  | Fullsib | 100 | | 99.5 | | 99.5 |
|  | Sire | 100 | | 97.8 | | 99.3 |
| 5% | Random | 100 | | 100 | | 99.9 |
|  | Fullsib | 100 | | 99.5 | | 99.9 |
|  | Sire | 100 | | 97.8 | | 99.9 |
| 10% | Random | 100 | | 100 | | 99.9 |
|  | Fullsib | 100 | | 99.5 | | 99.9 |
|  | Sire | 100 | | 97.9 | | 99.9 |

Here accuracy of pedigree correction was the proportion of the individuals whose sires were correctly identified

**Supplementary Table S2 accuracy of pedigree correction of genotypes with 1% of SNPs randomly sporadic missing and 0.5% of SNPs randomly containing error across all individuals (%)**

| Rate | Type | PEC | SeekParentF90 | | AlphaAssign | |
| --- | --- | --- | --- | --- | --- | --- |
| 1% | Random | 100 | | 100 | | 99.5 |
|  | Fullsib | 100 | | 100 | | 99.5 |
|  | Sire | 100 | | 99.3 | | 99.3 |
| 5% | Random | 100 | | 100 | | 99.9 |
|  | Fullsib | 100 | | 99.7 | | 99.9 |
|  | Sire | 100 | | 99.1 | | 99.9 |
| 10% | Random | 100 | | 100 | | 99.9 |
|  | Fullsib | 100 | | 99.8 | | 99.9 |
|  | Sire | 100 | | 99.0 | | 99.9 |

Here accuracy of pedigree correction was the proportion of the individuals whose sires were correctly identified
